# Transcription factor interactions promote binding to non-cognate DNA sites

**DOI:** 10.64898/2026.10.08.757710

**Authors:** Yongming James Du, Yinxia Yan, Mario Halic

## Abstract

Recruitment of transcription factors (TFs) to non-cognate DNA sequences is widely observed, but the underlying mechanisms remain poorly understood. Here, we show that, within the nucleosome, Sox2 can bind an Oct4 motif, interact with the Oct4 POU-specific domain, and displace the Oct4 homeodomain. We also observe Sox2 binding at multiple near-canonical and non-canonical DNA sites, with these interactions promoted by a second Sox2 molecule or by nearby nucleosome-bound Oct4. The TF–TF interactions can bridge across DNA gyres, bringing together binding sites that are distant in linear DNA but spatially proximal on the nucleosome. We further find that Sox2 can engage across nucleosomal DNA gyres through loop regions and/or intrinsically disordered segments, providing an additional mode of chromatin binding. Together, our structural and biochemical data reveal how TF–TF interactions change DNA recognition on the nucleosome, enabling binding to non-cognate sequences and expanding the repertoire of pioneer factor interactions on chromatin.

## Introduction

Transcription factors (TFs) frequently occupy genomic sites lacking their canonical DNA recognition motifs, as widely documented through genome-wide profiling techniques such as ChIP-seq ^1–4^. This observation challenges the simple, direct DNA-binding model and has led to different mechanistic hypothesis. One proposed mechanism involves tethered or piggyback binding, whereby a TF is recruited to non-motif sequences through direct physical interactions with a second DNA-bound TF, independently of the first TF’s DNA-binding domain ^5^. Pioneer TFs, which can engage nucleosomal DNA, further facilitate this process by opening chromatin and creating platforms that recruit follower TFs via cooperative protein-protein interactions (PPIs) or bridging factors ^6^.

Such hypotheses are supported by cases where deletion of a TF’s DNA-binding domain does not abolish its localization to target promoters, pointing to PPI-driven mechanisms ^7,8^. These interactions are evident in large-scale datasets from projects like ENCODE and are particularly relevant in developmental contexts, enhancer function, and cell fate specification. Understanding this layered recruitment expands models of transcriptional regulation beyond motif-centric views.

ChIP-seq datasets from embryonic stem cells reveal that a substantial fraction of Oct4- and Sox2-occupied sites lack strong canonical motifs for these transcription factors, suggesting that binding at these loci is supported by cooperative recruitment ^9^. Such co-recruitment can be mediated both through composite DNA elements and through direct protein–protein interactions. In cells the cooperative recruitment is essential for early development and efficient reprogramming ^10–16^, however, the molecular mechanisms underlying TF recruitment to non-cognate sequences remain incompletely understood.

Oct4 contains two distinct DNA-binding modules, the POU-specific (Oct4_POU_S_) and POU-homeodomain (Oct4_POUHD) domains. These domains wrap around naked DNA in a bipartite configuration ^17,18^; however, such binding mode is sterically incompatible with the nucleosome architecture. We previously demonstrated that Oct4 uses its acidic patch to engage histone tails, and that histone post-translational modifications can modulate the cooperative binding of Oct4 and Sox2 to nucleosome ^19^, however the mechanisms of this cooperative binding remained unclear.

To understand the molecular mechanisms underlying cooperative binding of Oct4 and Sox2 to chromatin, we determined multiple cryo–electron microscopy (cryo-EM) structures of human Oct4 and Sox2 bound to nucleosomes containing DNA sequences from the human *Lin28B* locus, complemented by biochemical assays. Our data reveal unexpectedly promiscuous Sox2 binding: Sox2 can engage multiple near-canonical motifs as well as several non-canonical motifs. Sox2 occupancy at non-canonical sites is promoted by protein-protein interactions with another Sox2 molecule or with Oct4 bound nearby, and its DNA binding mode is distinct from binding to canonical sites. Our structural and biochemical findings indicate that TF–TF interactions can remodel how these proteins engage DNA, enabling recruitment to non-canonical sequences and expanding the spectrum of chromatin-bound pioneer factor interactions.

## Results

### Oct4 and Sox2 binding to H3K27ac_Lin28B nucleosome

To analyze the co-recruitment of Oct4 and Sox2, we have assembled complex of Oct4 and Sox2 bound nucleosome reconstituted with a 182-bp DNA fragment from the human *LIN28B* locus (**Supplementary Information Fig. 1a-c**). This sequence has three Oct4-binding sites (OBS1–3, ATGTTAAT) and one predicted Sox2-binding motif (ACAATA), all of which are similar to canonical motifs (**Supplementary Information Fig. 1d, e**) *^9,19,20^*. We previously described how Oct4 binds to the unmodified Lin28B nucleosome at OBS1 in the linker region ^19^, and we defined this register as position 0. We have also shown in biochemical assays that H3K27ac promoted cooperative binding of Oct4 and Sox2 to the nucleosome by shifting the register by 1 or 2 bp *^19^*. Here, to determine the mechanism underlying the cooperativity between Oct4 and Sox2, we assembled H3K27ac-modified Lin28B nucleosomes with both TFs (**Supplementary Information Fig. 1a-c**) and collected cryo-EM data (**Supplementary Information Fig. 1f-g**).

**Figure 1.**
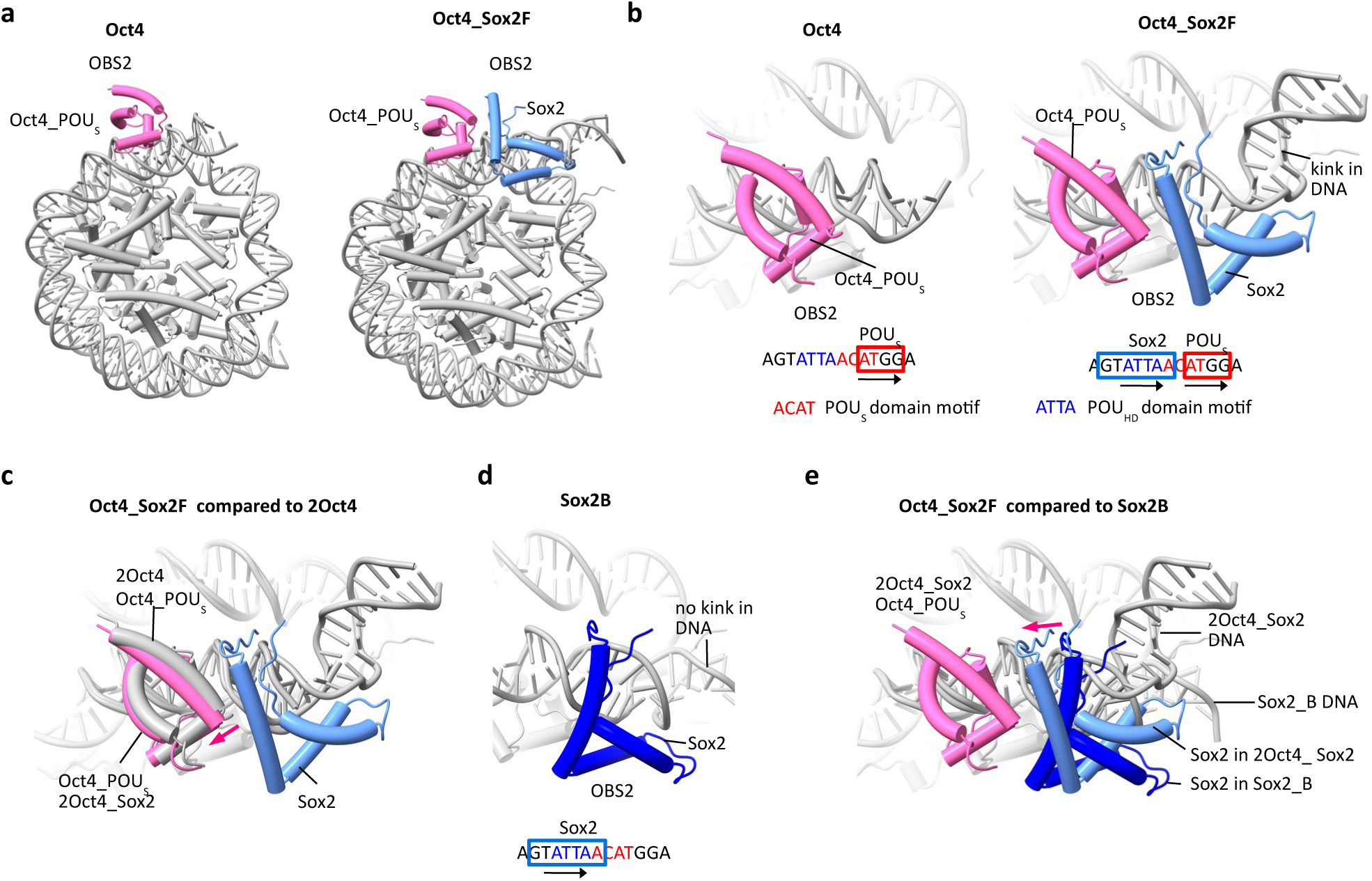
Sox2 binds Oct4 motif in context of nucleosome. **a,** Structural models of human Oct4 (Oct4, left) and Oct4:Sox2 (Oct4_Sox2F, right) bound to H3K27ac-modified nucleosomes (grey) assembled on a 182-bp DNA fragment from the *LIN28B* locus. **b,** Structural models of Oct4 (left) and Oct4:Sox2 (right) bound to OBS2 on H3K27ac nucleosomes. Bottom, OBS2 sequence is shown with Oct4 motif shown in blue and red. Binding sites for Oct4_POUS and Sox2 are shown as red and blue squares, respectively. **c,** Close-up view showing comparison of Oct4 binding to OBS2 in Oct4 structure (grey) with Oct4 binding in Oct4:Sox2F (pink) bound to OBS2 on H3K27ac nucleosomes. **d,** Structural model of Sox2 binding to OBS2 in Sox2B structure. Binding site for Sox2 is shown as blue square. **e,** Comparison of Sox2 binding to OBS2 in Oct4_Sox2F structure (light blue) with Sox2 binding in Sox2B (dark blue) bound to OBS2 on H3K27ac nucleosomes.

Classification of the data revealed 8 unique maps of Oct4 and/or Sox2 bound to H3K27ac nucleosomes (**Supplementary Information Fig. 1f, 2a, 2b and Table 1**). In all 8 maps, the nucleosome adopts position +1—a register shifted by 1 bp inward relative to the DNA positioning in the Oct4-bound unmodified nucleosome, consistently with our previous biochemistry observations (**Supplementary Information Fig. 3a, Table 2**). One map (referred to as Oct4) showed only Oct4 bound to the nucleosome at OBS2. Four maps showed 1 or 2 molecules of Sox2 bound at different positions (maps Sox2A, Sox2B, Sox2C, 2Sox2CD, wherein the letter marks a specific position for Sox2 and the number before the transcription factor indicates the number of molecules). Three maps showed both Oct4 and Sox2 bound, with different stoichiometries and at different positions (maps Oct4_Sox2E, Oct4_Sox2F, 2Oct4_2Sox2FG) (**Supplementary Information Fig. 1f and 2a-b, 4a-b**). Collectively in our structures, we see Oct4 bound at all of the 3 predicted sites (OBS1-3) (**Supplementary Information Fig. 1d**) ^19^.

**Figure 2.**
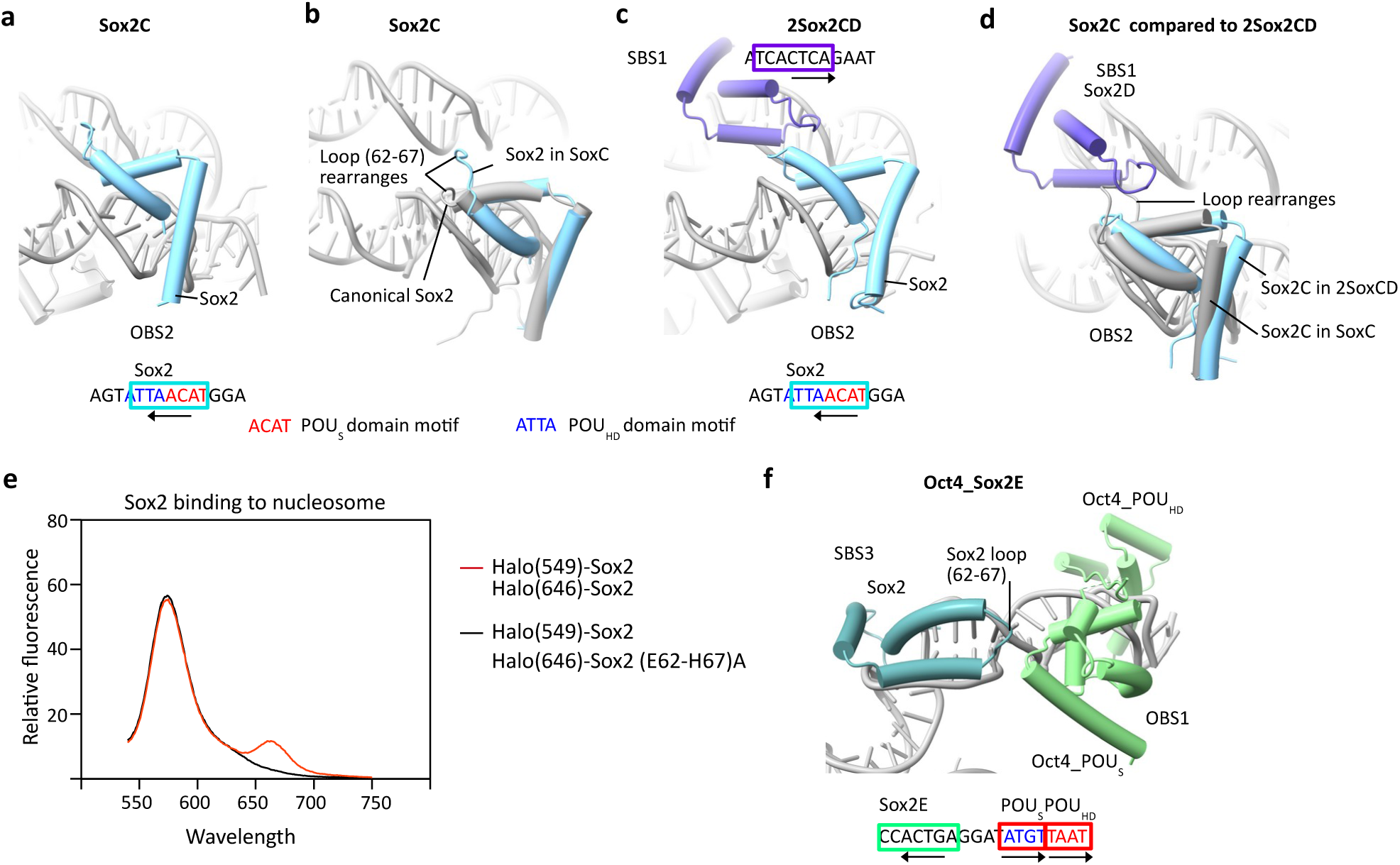
Oct4 and Sox2 recruit additional Sox2 molecules. **a,** Structural model of Sox2 binding to OBS2 in Sox2C structure. Binding site for Sox2 is shown as cyan square. **b,** Comparison of Sox2 binding to OBS2 in Sox2C structure (cyan) with canonical Sox2 binding (PDB:8bx2, grey). **c,** Structural model of Sox2 binding to OBS2 (cyan) and site D (purple) in Sox2CD structure. Binding site for Sox2 is shown as cyan and purple squares. **d,** Comparison of Sox2 binding to OBS2 in Sox2C structure (grey) and Sox2CD structure (cyan and purple). **e,** Emission spectra of Halo(549)_Sox2 and Halo(646)_Sox2 bound to nucleosome recorded upon donor excitation. Sensitized acceptor emission (660 nm) indicates FRET in the donor+acceptor condition. **f,** Structural model of Oct4 binding to OBS1 (light green) and Sox2 binding to site E (dark green) in Oct4_Sox2E structure. Binding sites for Oct4 and Sox2 are shown as red and green squares, respectively.

**Figure 3.**
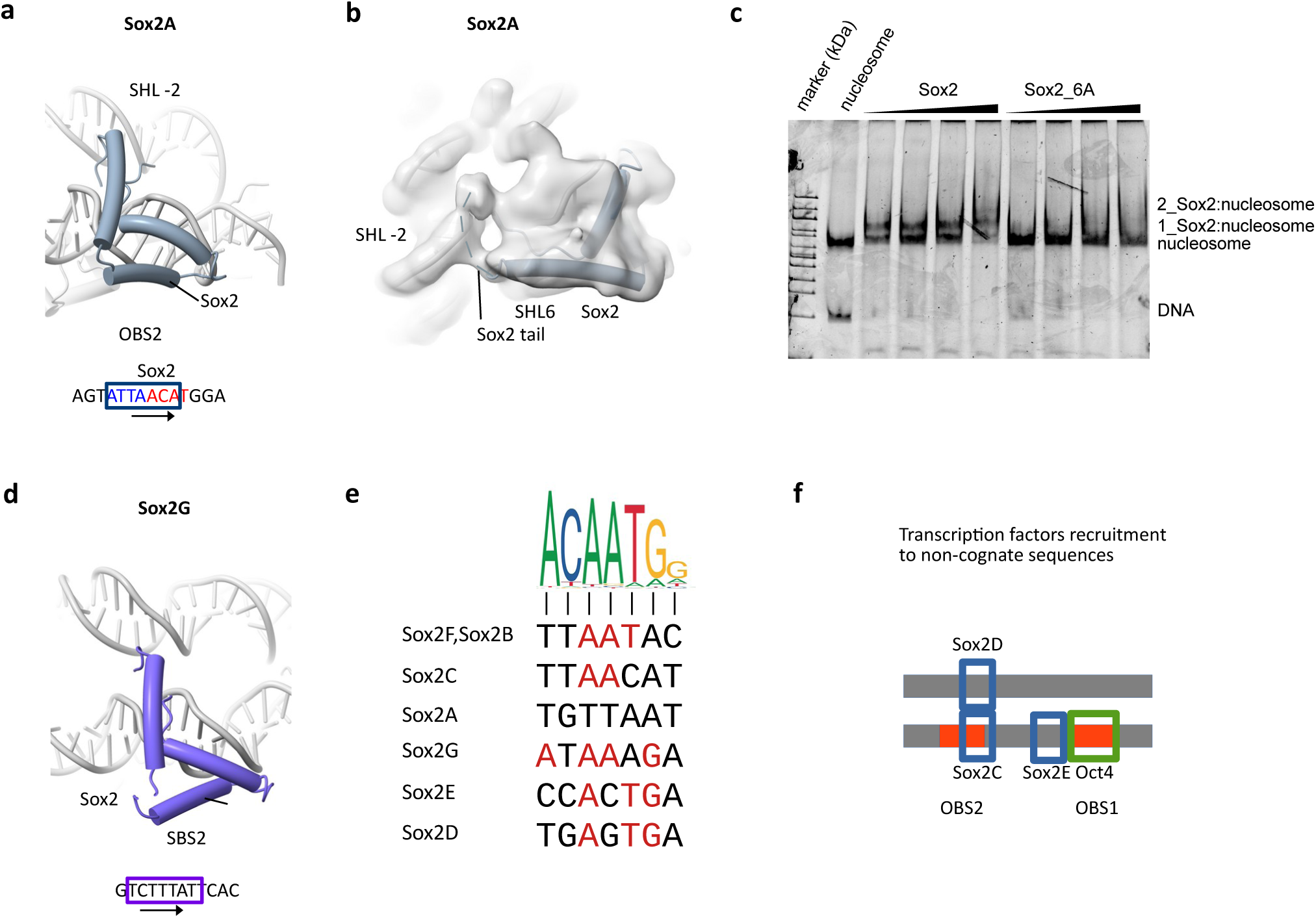
Oct4 and Sox2 recruit additional Sox2 molecules. **a,** Structural model of Sox2 binding to OBS2 in Sox2A structure. Binding site for Sox2 is shown as cyan square. **b,** Cryo EM density and structural model of Sox2 binding to OBS2 in Sox2A structure. Sox2 tail interacts with DNA across DNA gyres. **c,** Native gel showing binding of Sox2 and Sox2_6A to nucleosome. **d,** Structural model of Sox2 binding to site G (purple) in Sox2FG structure. Binding site for Sox2 is shown as purple square. **e,** Comparison of DNA sequences where Sox2 binds in our structure with canonical motif. Bases that are same as in canonical motif are colored red. **f,** Model showing transcription factor recruitment to non cognate sequences through protein-protein interactions.

We do not observe Sox2 bound to its predicted canonical motif, which is located at SHL4 in our nucleosome structures. We note that the minor groove at this position is occupied by the H2A N-terminal tail, which could sterically hinder Sox2 engagement (**Supplementary Information Fig. 5a**). These observations suggest that histone tails can compete with TFs for DNA access.

### Sox2 binds Oct4 motif

Instead of engaging with its predicted canonical site, we see that Sox2 engages with Oct4-binding site OBS2 (in maps Oct4_Sox2F, Sox2A, Sox2B, Sox2C) and multiple other non-canonical sites. Notably, Sox2 at OBS2 interacts with the canonical motif for Oct4 (**Fig. 1a, b**). In the Oct4_Sox2F map Sox2 is bound to TTAATCT, which is quite different from the Sox2 canonical motif ACAATGG, whereas the Oct4 POUS domain binds ATGG instead of the canonical ATGT which is also present at the site (**Fig. 1b**). Although the underlying sequences are different, the DNA binding mode of Oct4 and Sox2 in our Oct4_Sox2F map resembles that seen in previous structures of Oct4 and Sox2 bound to DNA (**Supplementary Information Fig. 5b**) ^18,21^.

We compared Oct4 binding to OBS2 in presence (Oct4_Sox2F) or absence of Sox2 (Oct4) (**Fig. 1c**). The comparison reveals that Sox2 rearranges the Oct4 molecule at OBS2, pushing Oct4 away, which alleviates the clash between them (**Fig 1c**). Moreover, co-binding of Sox2 and Oct4 at OBS2 induces a pronounced ∼90° kink in the DNA, which is not observed with Oct4 (Oct4) or with Sox2 (Sox2B) alone bound to nucleosome (**Fig. 1b, d**). In the latter, Sox2 binds the same sequence at OBS2 as in Oct4_Sox2F, indicating that simultaneous occupancy by both factors is required to drive this DNA conformational rearrangement (**Fig. 1d, e**).

As mentioned earlier, the DNA register in all our structures of the H3K27ac nucleosome with Oct4 and Sox2 is shifted by 1 bp relative to the structure of Oct4 bound to the unmodified nucleosome ^19^. To test whether this DNA repositioning could facilitate cooperative binding, we modeled the Oct4:Sox2 complex bound to OBS2 (Oct4_Sox2F) onto DNA in register 0 (**Supplementary Information Fig. 5c**). In this configuration, both Oct4 and Sox2 would clash with nucleosomal DNA and histones, indicating that cooperative binding requires local DNA unwrapping and/or DNA repositioning. These structural constraints provide a mechanistic rationale for how H3K27 acetylation can promote Oct4–Sox2 cooperativity ^19^, namely by increasing DNA accessibility and the register shift needed to accommodate the complex.

Together, these data show that Oct4 and Sox2 adopt distinct DNA-binding modes during cooperative engagement, and that co-binding of these two factors is necessary to generate a DNA kink. Such DNA deformation could weaken nucleosome–nucleosome contacts, promote local chromatin opening, and facilitate transcriptional activation.

### Sox2 and Oct4 recruit additional Sox2 molecules to non-canonical binding sites

In addition to Sox2 binding at OBS2 in the Oct4-associated context, we observe Sox2 engaging with other DNA sequences on the nucleosome. In the Sox2C structure, Sox2 binds the same site at OBS2 but in an inverted orientation (position C, **Fig. 2a**). Because TTAA is palindromic, the DNA readout is essentially the same, but the orientation of Sox2 in position C is incompatible with cooperative binding to Oct4 via the previously defined protein–protein interface (**Fig. 1b and Supplementary Fig. 5d**). Moreover, we see remodeling of Sox2 in position C: a loop (aa 62-67) adopts a distinct conformation, compared to Sox2 in Sox2B or Oct4_Sox2F, which enables contacts with DNA on the opposite gyre of the nucleosome relative to the canonical Sox2 binding mode (**Fig. 2b and Supplementary Fig. 5e**). Notably, the binding mode of Sox2 at position C in OBS2 allows it to engage a second Sox2 molecule, bound on the opposing DNA gyre near SHL2 (position D), as seen in the 2Sox2CD structure (**Fig. 2c**). To accommodate this Sox2–Sox2 interaction, both Sox2 at position C and the local DNA undergo rearrangements relative to the Sox2C structure (**Fig. 2d and Supplementary Fig. 5f**). A loop in Sox2 at position C remodels again to support the interaction with Sox2 at position D; the two Sox2 molecules contact one another across the nucleosomal DNA gyres primarily through the loop (aa 62-67) mediated interface.

In the 2Sox2CD structure, the Sox2 molecule at position D binds a non-canonical TGAGTGA motif at SHL2 and, unlike Sox2 at more canonical sites, does not induce the characteristic minor-groove widening that was seen in structures of Sox2 bound to canonical or near canonical motifs ^18^ (**Supplementary Fig. 5g**). This suggests that Sox2 at position D engages DNA in a non-canonical manner, with reduced sequence specificity; and is recruited or stabilized primarily through interactions with Sox2 at position C and with the DNA backbone.

To test whether Sox2 at position C recruits Sox2 at position D, we mutated loop residues 62–67, which form the Sox2–Sox2 interface, to alanine and fluorescently labeled Halo-tagged Sox2 with two dyes to enable FRET between Sox2 molecules. When bound to nucleosome, wild-type Sox2 exhibited a robust FRET signal, whereas the Sox2(62–67)A mutant showed no detectable FRET (**Fig. 2e**), indicating that this loop-mediated interaction is essential for recruitment of the second Sox2 to site D. Notably, FRET between Sox2 molecules was not observed on free DNA, in agreement with our structural work (**Supplementary Fig. 6a, b**). These data show that Sox2 can recruit additional Sox2 molecules to non-cognate DNA sites through direct protein–protein interactions.

In another map (Oct4_Sox2E), we observe Oct4 bound at OBS1 in the linker region interacting with a Sox2 molecule bound to another non-canonical motif (CCACTGA) (**Fig. 2f**). In this configuration, the same Sox2 loop (aa 62-67) contacts the Oct4 POUS domain, likely stabilizing Sox2 on this non-canonical sequence.

Together, these observations indicate that Sox2–Sox2 and Oct4–Sox2 interactions can recruit additional Sox2 molecules to non-canonical DNA sites, to which Sox2 binds in a sequence-independent way; and Sox2–nucleosome and Sox2–Sox2 interactions can drive conformational remodeling of both Sox2 and the nucleosomal DNA, expanding the repertoire of Sox2 binding geometries on chromatin.

### Diverse Sox2 binding modes on the nucleosome

In an additional set of structures, we observe further Sox2 binding configurations on the nucleosome. In the Sox2_A map, Sox2 engages OBS2 in an alternative binding mode and recognizes a TGTTAAT sequence, which corresponds to a near-canonical Sox motif in the reverse-complement orientation (**Fig. 3a**). This AT-rich motif facilitates minor-groove distortion and supports canonical Sox2 binding ^22,23^, in contrast to the binding modes observed at sites D and E, which depend on interactions with another transcription factor molecule. Notably, this binding does not allow for cooperative binding with Oct4. In this state, the Sox2 C-terminal tail extends across nucleosomal DNA gyres and contacts DNA, further expanding the repertoire of Sox2–nucleosome interactions, in agreement with recent report ^24^ (**Fig. 3b**). These data indicate that both the Sox2 DNA-binding loop and intrinsically disordered regions can contact nucleosomal DNA across gyres in a largely sequence-independent manner.

To test the importance of interactions between this intrinsically disordered region and DNA across nucleosomal gyres, we mutated residues R113/R114/K115/K121/K122/K124 in Sox2 disordered region to alanine, generating the Sox2_6A mutant. Sox2_6A bound naked DNA only slightly more weakly than wild-type Sox2 (**Supplementary Fig. 6c**); however, its binding to nucleosomes was strongly reduced (**Fig. 3c**). These data show that interactions between the charged intrinsically disordered region of Sox2 and nucleosomal DNA are essential for stable Sox2 binding to chromatin. In the 2Oct4_2Sox2FG map we identify an additional Sox2 molecule bound on the opposite side of the nucleosome near SHL −6, associated with yet another sequence (ATAAAGA) (**Fig. 3d**). In summary, our data show that Sox2 can bind a diverse range of sequences and locations on the nucleosome (**Fig. 3e**). The AT-rich core is sufficient to support Sox2 binding (sites A, B, C, G), whereas at sites lacking this feature (sites D, E), Sox2 recruitment is assisted by interactions with a nearby Oct4 or Sox2 molecule.

## Discussion

Taken together, our structures reveal unexpected complexity in how Oct4 and Sox2 engage nucleosomal DNA, extending the classical view of Oct4–Sox2 cooperativity derived from naked-DNA complexes. Oct4 and Sox2 were originally established as core pluripotency factors that act cooperatively at composite cis-regulatory elements, where direct protein–protein interactions and motif spacing/orientation are critical for enhancer activation ^18,25^. More recent work has emphasized that pioneer-factor function must be understood in the context of chromatin rather than naked DNA, because nucleosome geometry can strongly reshape motif accessibility, factor conformation, and cooperative assembly ^6,9,19,26–29^. In this context, our cryo-EM analysis shows that Oct4 and Sox2 do not simply recognize pre-encoded composite motifs on nucleosomal DNA. We show that Sox2 can occupy the Oct4 motif by binding the motif region normally engaged by the Oct4 POU_HD_ domain. This shows that Oct4 and Sox2 exploit chromatin-dependent structural plasticity, adopting multiple binding modes that depend on local DNA register, histone contacts, and the presence of partner factors. These observations support a model in which nucleosomes act not only as barriers to transcription factor binding, but also as allosteric scaffolds that redirect specificity and cooperativity ^30^.

Our data also show that cooperative Oct4–Sox2 binding at OBS2 drives a pronounced ∼90° DNA kink. The fact that Oct4 and Sox2 alone can bind the same sequence without producing the kink indicates that the deformation is an emergent property of the cooperative complex rather than an intrinsic feature of Sox2 engagement at that site. Such localized DNA distortion could weaken nearby histone– DNA or nucleosome–nucleosome contacts and thereby contribute to chromatin opening at pluripotency enhancers, consistent with the long-standing view that pioneer factors initiate transcriptional activation by remodeling otherwise inaccessible chromatin ^26^.

Nucleosome-bound Sox2 is conformationally adaptive rather than operating as a rigid reader of a single consensus motif. Sox2 appears to remodel its own loops and disordered regions in response to the nucleosomal surface. This is consistent with data showing that transcription factors can display altered recognition modes on nucleosomes compared with naked DNA, because the histone octamer imposes asymmetric groove exposure, DNA pre-bending, and local steric restrictions ^9^.

Our structures also highlight an underappreciated role for histone tails in shaping transcription factor occupancy. Despite the presence of a predicted canonical Sox2 site at SHL4, we do not observe Sox2 bound there; instead, the minor groove at that position is occupied by the H2A N-terminal tail. Histone tails are often discussed as substrates for post-translational modification and recruiters of chromatin effectors, but they also make extensive contacts with DNA and can directly modulate DNA accessibility on the nucleosome surface ^31^. Our data support the idea that histone tails can physically compete with transcription factors for access to DNA grooves. In this view, histone tail acetylation may facilitate factor binding also by displacing tail segments from DNA surfaces. This provides a plausible mechanistic bridge between histone modification and pioneer-factor access at individual nucleosomal sites.

A recent study showed that transcription factors frequently interact on DNA to form DNA-guided cooperative complexes that recognize composite motifs with defined spacing and orientation ^32^. Our work extends this concept to chromatin by showing that TF–TF interactions can occur on the nucleosome, including across DNA gyres, where DNA wrapping brings distant sites into proximity and enables binding to near-canonical or non-canonical motifs. Whereas the CAP-SELEX study defines sequence rules for TF cooperativity on free DNA, our structural and biochemical data reveal how TF– TF interactions reshape DNA recognition in a nucleosomal context.

The ability of Oct4–Sox2 and Sox2–Sox2 interactions to recruit additional Sox2 molecules to non-canonical sites may have important implications for enhancer assembly in vivo (**Fig 3f**). Regulatory elements occupied by pluripotency factors often contain clusters of degenerate motifs and can form cooperative assemblies that are difficult to explain by one-factor/one-site models ^33,34^. Our structures suggest that once an initial high-affinity or structurally favorable interaction is established, partner-factor contacts and nucleosome-assisted conformational remodeling can stabilize additional low-specificity binding events on the same particle. Such multivalent assemblies could help explain how Oct4 and Sox2 nucleate enhancer activation, recruit cofactors, and promote local chromatin accessibility even at sites that lack ideal consensus motifs. More broadly, these data align with emerging views that enhancer function depends on combinatorial and dynamic assemblies whose stability arises from many individually weak interactions rather than a single high-specificity recognition event ^35^.

## Methods

### Cloning and mutagenesis

The coding sequences for full-length and short form (residues 40-156) human Sox2 were amplified by PCR and cloned into the pETDuet-1 vector between the *EcoR*I and *Not*I restriction sites. For the generation of Halo-tagged Sox2 short form, the DNA fragment encoding Sox2 (residues 40-156) with an N-terminal GGGGS linker was inserted into the Halo-tag vector (Addgene, #188943), which was linearized via PCR amplification followed by *Dpn*I digestion to remove the parental template. The insertion was performed using Sequence-and-Ligation-Independent Cloning (SLIC). Two distinct mutants were generated via inverse PCR using Phusion Flash High-Fidelity PCR Master Mix (Thermo Scientific, F548) and mutagenic primers: the IDR R/K region mutant, in which multiple alanine (Ala) substitutions at residues 113-115, 121-122 and 124 were introduced sequentially, and the HMG loop1 mutant, featuring the simultaneous alanine substitution of consecutive residues 63-68. PCR amplification was followed by *Dpn*I digestion of the parental template and plasmid recirculation. All oligonucleotides used for cloning and mutagenesis are listed in **Supplementary Table 3**. All constructs and mutations were verified by Sanger sequencing. All plasmids used in this study are listed in **Supplementary Table 4**.

### Histone expression and purification

*Xenopus laevis* histones (H2A, H2B, H3, and H4) were overexpressed separately in BL21(DE3) pLysS cells and purified as described previously ^36–38^. Briefly, cells were grown to OD_600_ = 0.6 and induced with 1 mM IPTG for 3 h at 37 °C. After sonication and centrifugation, the inclusion bodies were pelleted and were solubilized for 1 h at room temperature in denaturing buffer with 50 mM Tris–HCl pH 7.5, 2 M NaCl, 6 M guanidine hydrochloride, 1 mM DTT. Equimolar H2A and H2B (for H2 dimer) or H3 and H4 (for H3/H4 tetramer) were mixed for 1 h at room temperature followed by successive overnight dialyses at 4 °C against 1 L refolding buffer that contains 25 mM HEPES–NaOH pH 7.5, 2 M NaCl, 1 mM DTT. The H2A/H2B dimer and H3/H4 tetramer were further purified by DEAE and SP resin and then size-exclusion chromatography with Superdex 200 column. SEC peak fractions in were pooled and concentrated for histone octamer assembly.

### Lysine-27 acetylated histone H3 expression and purification

H3K27ac modified histone H3 was prepared same as described ^39^. Briefly, the PyIT–H3 plasmid and pBK-AcKRS3 were co-transformed into BL21(DE3) pLysS cells. Selected colony on LB agar containing kanamycin and spectinomycin were tested for expression and was used for scale-up expression. For large-scale production, 8 L cultures were first grown at 37 °C to OD_600_ = 0.8, and then supplemented with 20 mM nicotinamide and 10 mM acetyl-lysine, 30 min later induced with 0.5 mM IPTG for 3 h at 37 °C. The harvested cells were washed with PBS containing 20 mM nicotinamide, and resuspended in 50 mL lysis buffer containing 50 mM Tris–HCl pH 7.5, 150 mM NaCl, 1 mM DTT, 1% Triton X-100. Cells were lysed by sonication on ice, and the lysate was clarified by centrifugation 37,000 × g for 20 min at 4 °C. The pellet was solubilized in 25 mL denaturing buffer with 20 mM Tris– HCl pH 8.0, 6 M guanidine HCl and incubated 2 h at room temperature with gentle agitation, and clarified again (37,000 × g, 15 min, 4 °C).

The supernatant was purified by 5 mL Ni-NTA resin pre-equilibrated in 20 mM Tris–HCl pH 8.0, 0.5 M NaCl, 6 M urea, and incubated 1 h at 4 °C. The resin was washed by 50 mL equilibration buffer and then 100 mL the equilibration buffer supplemented with 20 mM imidazole and eluted with 10 mL elution buffer (equilibration buffer plus 300 mM imidazole). The H3K27ac/H4 tetramer was then prepared by dialysis using the same procedure as for the unmodified H3/H4 tetramer, and concentrated to ∼1 mg/mL.

### Histone octamer preparation

Two-fold molar excess of H2A/H2B dimer was mixed with H3/H4 tetramer containing either unmodified or H3K27ac-modified H3 and incubated for 1 hon ice. Histone octamer was purified by size-exclusion chromatography using Superdex 200 Increase 10/300 GL column in buffer with 25 mM HEPES–NaOH (pH 7.5), 2 M NaCl, 1 mM DTT. The peak fractions were confirmed by SDS–PAGE concentrated, and aliquoted for nucleosome assembly.

### Nucleosome DNA preparation

Human LIN28B 182-bp DNA was prepared as described previously ^19^. PCR-amplified DNA were purified by phenol–chloroform extraction followed by ethanol precipitation. DNA integrity and purity were assessed by native PAGE gel.

Human LIN28B 182bp DNA sequence: gcataagttaagtggtattaacatatcctcagtggtgagtattaacatggaacttactccaacaatacagatgctgaataaatgtagtctaagtgaaggaagaaggaaaggtgggagctgccatcactcagaattgtccagcagggattgtgcaagcttgtgaataaagacacatacttcat

### Nucleosome assembly

Both unmodified and H3K27ac nucleosomes were reconstituted with same dialysis procedure as previously discribed ^36,40^. Briefly, the 182 bp LIN28B DNA fragment and purified histone octamer were mixed in a 200 µL dialysis button with high salt assembly buffer (25 mM Tris–HCl pH 7.5, 2 M NaCl, 1 mM DTT), placed inside a 6-8 kDa MWCO dialysis bag containing 50 mL of the same buffer, stirred in 1 L of buffer (25 mM Tris–HCl pH 7.5, 1 M NaCl, 1 mM DTT) for overnight dialysis at 4 °C. The bag was then moved to 1 L of low-salt buffer (25 mM Tris–HCl pH 7.5, 1 mM DTT) and dialyzed for an additional 6 h at 4°C. Reconstituted nucleosomes were concentrated to 2-5 µM by a 0.5 mL Amicon centrifugal filter (50 kDa MWCO). Nucleosome quality was assessed by 6% native PAGE run in 0.5× TBE.

### Purification and refolding of His tagged human Oct4 and Sox2 full-length protein

Full-length human Oct4 and Sox2 with an N-terminal His tag was expressed in BL21(DE3) pLysS cells using the same induction conditions as for histone expression. Cells were lysed by sonication in lysis buffer (50 mM Tris–HCl pH 7.5, 150 mM NaCl, 1% Triton X-100, 1 mM DTT). Inclusion bodies were pelleted by centrifugation at 37,000 × g for 15 min at 4 °C. The pellet was solubilized in 25 mL denaturing buffer (20 mM Tris–HCl pH 8.0, 6 M guanidine hydrochloride) and incubated for 2 h at room temperature with gentle agitation, followed by clarification at 37,000 × g for 15 min at 4 °C. The supernatant was purified by 5 mL Ni resin following the procedure used for histone puriffication. For refolding, purified Oct4 was dialyzed overnight at 4 °C against 200 mL buffer (25 mM HEPES–NaOH pH 7.5, 2 M urea, 250 mM NaCl, 1 mM DTT), then dialyzed for an additional 6 h at 4 °C against 200 ml of 25 mM HEPES–NaOH pH 7.5, 100 mM NaCl, 1 mM DTT. The dialyzed sample was clarified by centrifugation at 21,300 × g for 15 min at 4 °C. Refolding and activity were verified by SDS–PAGE and electrophoretic mobility-shift assays (EMSA) using LIN28B DNA to confirm DNA-binding activity.

### Oct4/Sox2-Lin28B H3K27ac nucleosome complex preparation for cryo-EM analysis

For preparation Oct4/Sox2-Lin28B H3K27ac nucleosome complex, Lin28B H3K27ac nucleosome(5 mM), refolded Oct4, refolded Sox2 (molar ratio: 1/2/4) were mixed in 500 mL TND buffer and incubated at 25°C for 15 minutes at and concentrated to 1 mg/mL for grid prep using a 100 kDa MWCO Amicon filter.

### Cryo-EM grid preparation and data collection

Nucleosome complex sample was kept on ice after concentration and directly loaded 3 mL onto the glow-discharged holey carbon grid (Quantifoil 300 mesh Cu R1.2/1.3). The humidity in the chamber was kept at 95% and the temperature at 10°C. After 3 second blotting, grids were plunge-frozen in liquid ethane using Thermo Scientific Mark IV vitrobot system. Electron micrographs were recorded on 300k eV Titan Krios electron microscope with a K3 electron detector at the Cryo-EM facility at St. Jude Childrens’s Research Hospital. Image pixel size was 0.6485 Å per pixel on the object scale. Data were collected in a defocus range of 9,000–30,000 Å with a total exposure of 80 e^-^Å^-2^. Sixty frames were collected.

Frames were aligned with the MotionCorr2 software. The contrast transfer function parameters were determined using CTFFIND4. Particles were picked by Topaz using pre-trained model. Particles were windowed and 2D class averages were generated with the Relion software package ^41,42^. Inconsistent class averages were removed from further data analysis. The initial reference was filtered to 40 Å in Relion. C1 symmetry was applied during refinements for all classes. Particles were split into 2 datasets and refined independently and the resolution was determined using the 0.143 cut-off (Relion auto refine option). All maps were filtered to resolution using Relion with a B-factor determined by Relion. High resolution maps were modified with EMReady ^43^.

### Model building and refinement

Molecular models were built in UCSF ChimeraX ^44^ and Coot ^45^. Oct4 bound H3K27ac LIN28B nucleoosme complex was generated by rigid-body fitting of our previously determined Oct4–LIN28B nucleosome complex (PDB: 8G8G) structure. For the Sox2 alone and Oct4-Sox2 co-bound H3K27ac LIN28B nucleoosme complex, Oct4–LIN28B nucleosome complex (PDB: 8G8G) and Oct4/Sox2-nucleosome complex structure (PDB: 6T90) were used as initial models in model building. The LIN28B nucleosome and Oct4, Sox2 from initial models were fit independently into the cryo-EM map density in ChimeraX, manually adjusted in Coot, and refined by Phenix real-space refinement. Map– model correlation coefficients (CC) for DNA bases used to assess nucleosomal DNA register were computed with Phenix “Comprehensive validation (cryo-EM),” using models containing only nucleotide bases and the core nucleosome map. All cryo-EM maps and structural models were visualized in ChimeraX.

### Expression and Purification of His- and Halo-tagged Sox2_S WT and mutant proteins

LOBSTR *E. coli* were grown in 1 L LB at 37 °C, 220 rpm to OD600 ≈ 0.6, shifted to 16 °C and cultured to OD600 ≈ 0.8, then induced with 0.3 mM IPTG for 16 h at 16 °C, 180 rpm. Cells were harvested, washed with 50 mL buffer A (50 mM Tris–HCl pH 7.5, 150 mM NaCl), resuspended in buffer B (20 mM HEPES–NaOH pH 7.5, 1 M NaCl) supplemented with 1 mM PMSF, 1 mM DTT, and 0.1 mg/mL DNase I, and lysed by sonication. The clarified supernatant was incubated with 5 mL Ni-NTA resin for 1 h at 4 °C, washed with buffer B containing 30 mM imidazole, and eluted with buffer B containing 300 mM imidazole. The His-affinity–purified protein was further purified by size-exclusion chromatography in 20 mM HEPES–NaOH pH 7.5, 0.5 M NaCl, 1 mM DTT. Peak fractions were assessed by SDS–PAGE; intact full-length protein was pooled and concentrated to 5 mg/mL.

### Labeling of Halo-tag Sox2 proteins by Janelia Fluor fluorescence ligand

For fluorescence labeling, purified Halo tagged Sox2 proteins were incubated with 1.2 molecular ratio of Janelia Fluor fluorescence ligand Halo ligand-JF549 (Promega, GA1111) or JF646 (GA1121) overnight in 20 mM HEPES–NaOH pH 7.5, 150 mM NaCl, 1 mM DTT. The successfully labeled protein was purified by Superdex 200 Increase 10/300 GL column using the same buffer.

### EMSA nucleosome binding assay

To test binding between LIN28B nucleosome and Sox2 alone or Oct4 plus Sox2, 20 nM nucleosome or free DNA (reference) in buffer containing 20 mM Tris–HCl pH 7.5, 100 mM KCl, 1% glycerol, 0.2 mM EDTA, 0.2% CHAPS, 1 mM DTT was incubated with Oct4 or Sox2 proteins in the indicated ratios for 15 min, at 25 °C. The binding was visualized by electrophoresis in 6% native PAGE gel, 200V 30 min using 0.5x TBE buffer.

### FRET

To measure the fluorescence resonance energy transfer (FRET), 40 nM LIN28B nucleosome or free DNA in binding buffer (20 mM Tris–HCl pH 7.5, 100 mM KCl, 1% glycerol, 0.2 mM EDTA, 0.2% CHAPS, 1 mM DTT) was mixed with Janelia Fluor 549 (JF549) labeled Halo-Sox2 WT protein with molar ratio 1:2 and incubated at 25°C for 15 min, and incubated with 8 molar ratio JF646 Halo-Sox2 WT or Sox2(62-67)A mutant protein for another 15 min. The FRET was measured by FlexStation 3 (Molecular Devices) microplate reader with Excitation wavelength 500 nm and record the emission spectrum between 540 nm and 750 nm per 1 nm. Three replicates were performed for each binding.

## Acknowledgments

We thank Cryo-EM Center members at St. Jude Children’s Research Hospital for technical assistance and support in cryo-EM facility. We thank professor Jason Chin at University of Cambridge for sharing plasmids for acetylated histone preparation. We thank Hai Dao and Ashish Verma for providing initial H3K27ac histones. We thank Inês Chen for critical reading and comments. Work in the Halic laboratory is funded by St. Jude Children’s Research Hospital, the American Lebanese Syrian Associated Charities and NIH award R35GM158165. James Du was also supported by a Gephardt Postdoctoral Fellowship from the Cancer Biology Program, Comprehensive Cancer Center.

## Author contribution

Y.D. and M.H. conceived the study. Y.D. performed the experiments, with the exception of cloning, purification of Halo-Sox2, and FRET measurements, which were performed by Y.Y. Y.D. and M.H. wrote the manuscript with input and comments from all other authors.

## Data availability

EM density maps and models have been deposited in the Electron Microscopy Data Bank and PDB under the following accession codes: for Oct4 bound to H3K27ac Lin28B nucleosome, PDB xxxx was built using map EMD-xxx (), PDB xxxx was built using map EMD-xxx (), PDB xxxx was built using map EMD-xxx (), PDB xxxx was built using map EMD-xxx (), PDB xxxx was built using map EMD-xxx (). All other data supporting the findings of this study are available within the article and its supplementary information files.

## Supplementary Information

**Supplementary Figure 1.**
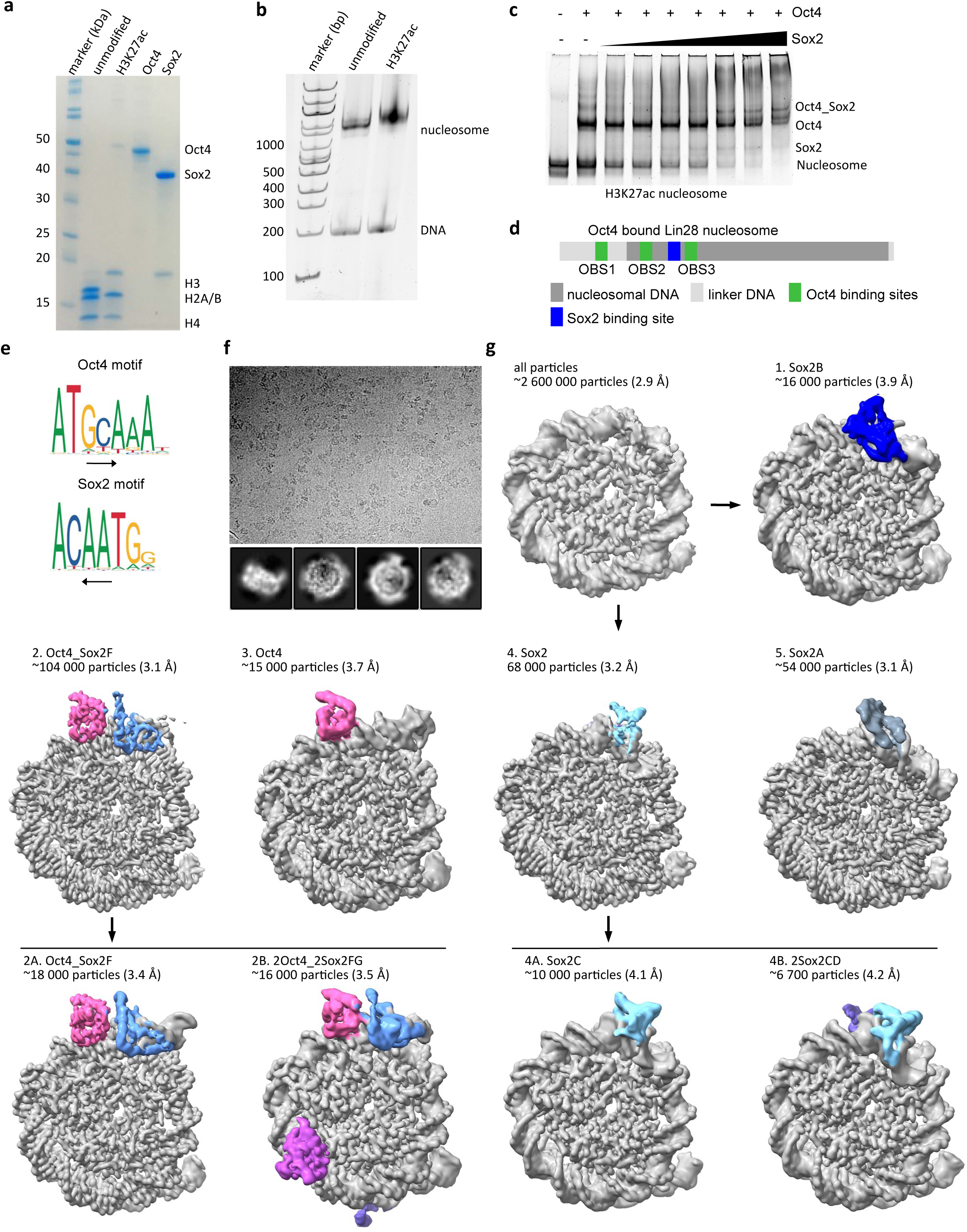
Assembly and cryo-EM of Sox2 and Oct4 bound to H3K27ac *LIN28B* nucleosomes. **a,** A SDS gel showing purification of Oct4, Sox2 and assembly of unmodified and H3K27ac histone octamer. **b,** Representative native gel electrophoresis showing assembly of unmodified or H3K27ac *LIN28B*nucleosomes. **c,** Representative native gel electrophoresis showing Oct4 and Sox2 binding to the H3K27ac *LIN28B* nucleosomes. **d,** Schematic representation showing *LIN28B* DNA positioning on the Oct4 bound nucleosome complex. Oct4 and predicted Sox2 binding sites are labeled. **e,** Canonical motifs for Oct4 and Sox2 (Jaspar). **f,** Top. Representative cryo-EM micrograph from a set of 78 069 micrographs collected with Titan Krios electron microscope at 300 keV. Nucleosome particles in multiple orientations are visible. Bottom, representative 2D class averages showing nucleosomes. **g,** Cryo-EM map of nucleosome from the entire dataset, refined to 2.9 Å. Classification of these data resulted in 8 major classes of Oct4 and Sox2 bound nucleosomes. Number of particles and resolution corresponding to each class is indicated.

**Supplementary Figure 2.**
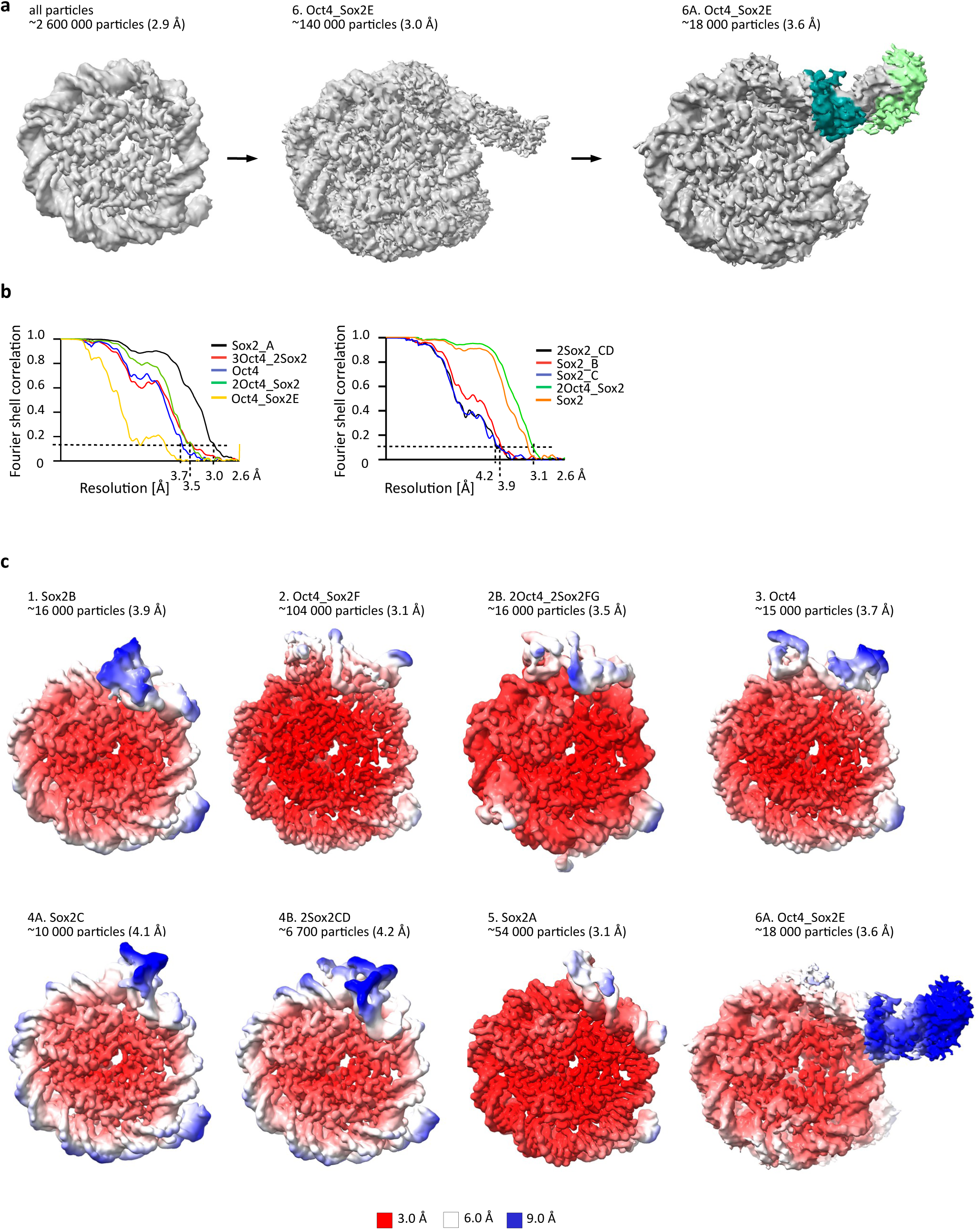
Cryo-EM of Oct4 and Sox2 bound to H3K27ac *LIN28B* nucleosomes. **a,** Further classification of data from S1f resulted in a class of Oct4 and Sox2 bound nucleosomes. Number of particles and resolution corresponding to each class is indicated. **b,** Fourier shell correlation (FSC) curve showing the resolution of maps from S1e and S2a. **c,** Cryo-EM maps shown in Figure 1f and 2a colored by local resolution.

**Supplementary Figure 3.**
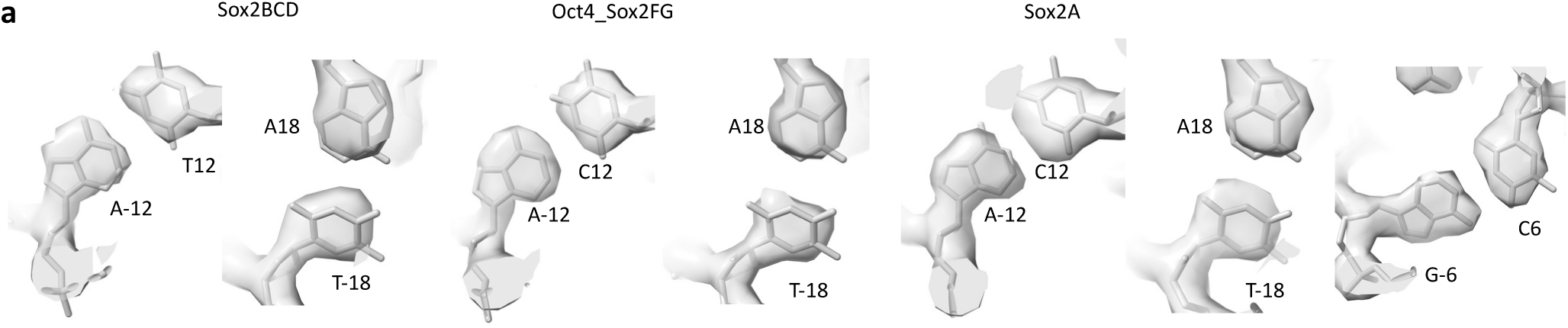
Model building of Oct4 and Sox2 bound to H3K27ac *LIN28B*nucleosomes. **a,** The representative region showing map quality of nucleotides and fit of corresponding models is shown for the nucleosome of indicated maps.

**Supplementary Figure 4.**
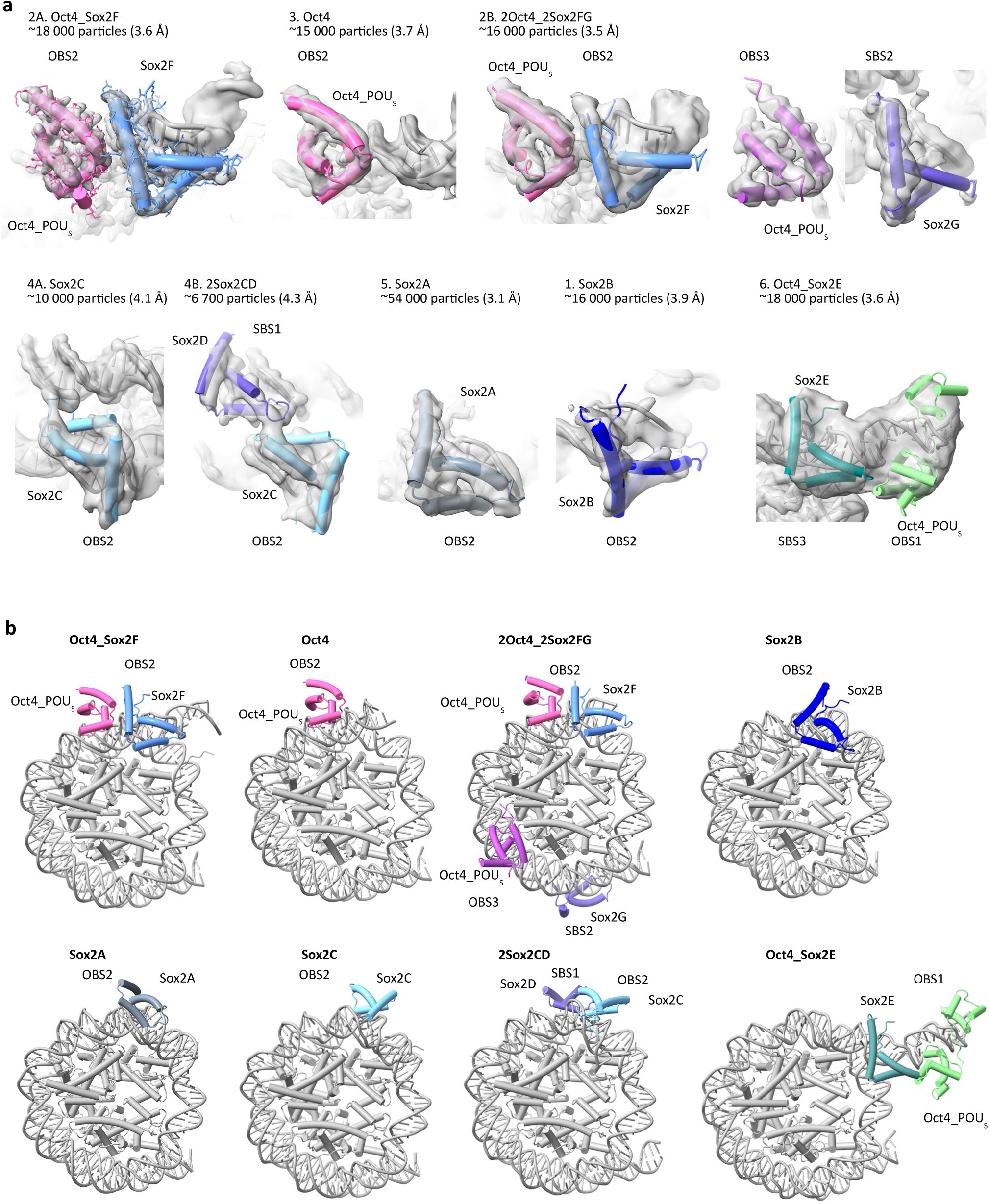
Model building of Oct4 and Sox2 bound to H3K27ac *LIN28B*nucleosomes. **a,** Cryo-EM maps of Oct4 and Sox2 bound to H3K27ac LIN28B nucleosome with fitted model showing Oct4 and Sox2 binding. **b,** Models of Oct4 and Sox2 binding to H3K27ac LIN28B nucleosomes.

**Supplementary Figure 5.**
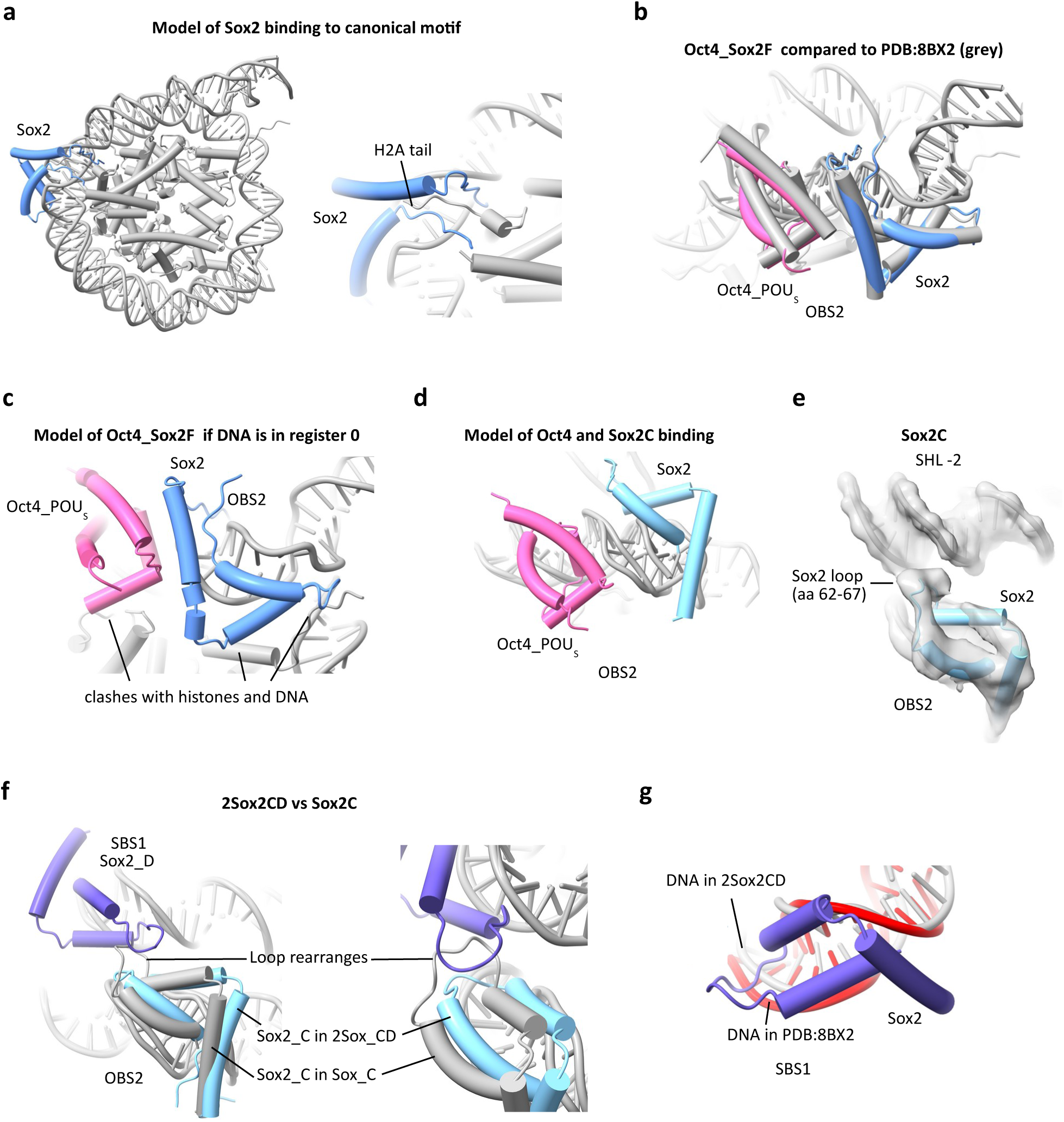
Cryo-EM of Oct4 and Sox2 bound to H3K27ac *LIN28B* nucleosomes. **a,** Model of Sox2 binding to predicted canonical motif. **b**, Comparison of Oct4 (pink) and Sox2 (blue) binding to OBS2 (2Oct4_Sox2F map) (grey) with Oct4 and Sox2 binding in PDB:8BX2. **c,** Predicted model of Oct4 and Sox2 binding to OBS2 when DNA is in register 0. Note clash with histones, which would not allow binding to this place. **d,** Predicted model of Oct4 and Sox2 binding to OBS2 when Sox2 is bound as in Sox2C structure. **e,** Cryo-EM map of Sox2C bound to H3K27ac LIN28B nucleosome with fitted model showing Sox2 loop binding DNA across DNA gyres in nucleosome. **f**, Comparison of Sox2 bound to site C in Sox2C (gray) and Sox2CD (light blue) structures showing rearrangement of Sox2. **g**, Comparison of the Sox2 bound DNA minor groove at site D with Sox2 bound to a canonical motif. Binding to the non-canonical site D site does not induce the characteristic minor-groove rearangement.

**Supplementary Figure 6.**
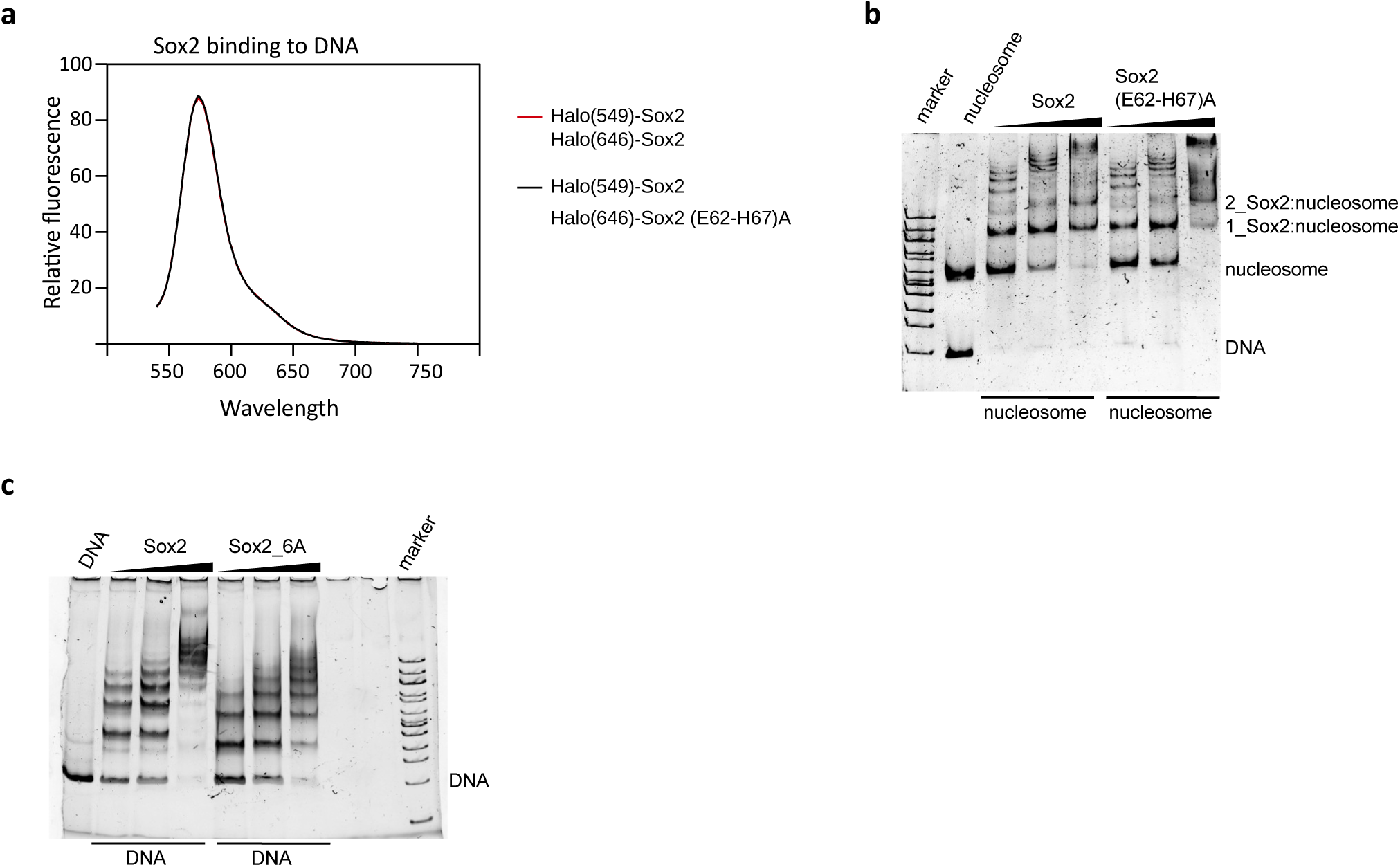
Sox2 recruits additional Sox2 molecule to nucleosome. **a,** Emission spectra of Halo(549)_Sox2 and Halo(646)_Sox2 bound to DNA recorded upon donor excitation. When bound to DNA FRET between two Sox2 molecules was not observed. **b,** Native gel showing binding of Sox2 and Sox2(E62-H67)A to nucleosome. **c,** Native gel showing binding of Sox2 and Sox2_6A to DNA.

**Supplementary Table 1.** Cryo-EM statistics.

|  | Sox2A | Sox2B | Sox2C | Sox2CD | Oct4 |
| --- | --- | --- | --- | --- | --- |
|  | EMD-xx | EMD-xx | EMD-xx | EMD-xx | EMD-xx |
|  | PDB ID xx | PDB ID xx | PDB ID xx | PDB ID xx | PDB ID xx |
| **Data collection and processing** |  |  |  |  |  |
| Magnification | 130,000 | 130,000 | 130,000 | 130,000 | 130,000 |
| Voltage (kV) | 300 | 300 | 300 | 300 | 300 |
| Electron exposure (e-/Å<sup>2</sup>) | 80 | 80 | 80 | 80 | 80 |
| Defocus range (μm) | -0.9 – -3.0 | -0.9 – -3.0 | -0.9 – -3.0 | -0.9 – -3.0 | -0.9 – -3.0 |
| Number of frames | 60 | 60 | 60 | 60 | 60 |
| Pixel size (Å) | 0.6485 | 0.6485 | 0.6485 | 0.6485 | 0.6485 |
| Symmetry imposed | C1 | C1 | C1 | C1 | C1 |
| Initial particle images (no.) | 2 600 000 | 2 600 000 | 2 600 000 | 2 600 000 | 2 600 000 |
| Final particle images (no.) | 54 000 | 16 000 | 10 000 | 6 700 | 15,000 |
| Map resolution (Å) | 3.1 | 3.9 | 4.1 | 4.2 | 3.7 |
| FSC threshold |  |  |  |  |  |
| Map resolution range (Å) | 2.9-9 | 3.7-9 | 3.8-9 | 3.8-9 | 3.5-6 |
| **Refinement** |  |  |  |  |  |
| Initial model used | 8g8g, 6t90 | 8g8g, 6t90 | 8g8g, 6t90 | 8g8g, 6t90 | 8g8g |
| Model resolution (Å) | 3.1 | 3.6 | 3.7 | 3.9 | 3.2 |
| FSC threshold |  |  |  |  |  |
| Model resolution range (Å) | 3.1-124 | 3.6-124 | 3.7-124 | 3.9-124 | 3.2-124 |
| **Model composition** |  |  |  |  |  |
| Nonhydrogen atoms | 12649 | 12649 | 12530 | 13017 | 12270 |
| Protein residues | 840 | 840 | 829 | 886 | 826 |
| Nucleotide | 290 | 290 | 289 | 288 | 278 |
| **B factors (Å<sup>2</sup>)** |  |  |  |  |  |
| Protein | 92.49 | 135.4 | 142.92 | 132.62 | 125.6 |
| Nucleotide | 134.3 | 237.36 | 238.4 | 245.52 | 147.52 |
| **R.m.s. deviations** |  |  |  |  |  |
| Bond lengths (Å) | 0.004 | 0.004 | 0.007 | 0.005 | 0.004 |
| Bond angles (°) | 0.64 | 0.76 | 0.839 | 0.952 | 0.755 |
| **Validation** |  |  |  |  |  |
| MolProbity score | 1.56 | 2.01 | 2.58 | 2.63 | 2.15 |
| Clashscore | 6.63 | 13 | 19.64 | 24.1 | 11.46 |
| Poor rotamers (%) | 1.57 | 2.28 | 3.91 | 3.62 | 2.89 |
| **Ramachandran plot** |  |  |  |  |  |
| Favored (%) | 97.8 | 97.44 | 95.07 | 95.27 | 96.53 |
| Allowed (%) | 2.20 | 2.56 | 4.93 | 4.62 | 3.47 |

|  | Oct4_Sox2E | 2Oct4_Sox2F | 3Oct4_Sox2FG |
| --- | --- | --- | --- |
|  | EMD-xx | EMD-xx | EMD-xx |
|  | PDB ID xx | PDB ID xx | PDB ID xx |
| **Data collection and processing** |  |  |  |
| Magnification | 130,000 | 130,000 | 130,000 |
| Voltage (kV) | 300 | 300 | 300 |
| Electron exposure (e-/Å<sup>2</sup>) | 80 | 80 | 80 |
| Defocus range (µm) | -0.9 – -3.0 | -0.9 – -3.0 | -0.9 – -3.0 |
| Number of frames | 60 | 60 | 60 |
| Pixel size (Å) | 0.6485 | 0.6485 | 0.6485 |
| Symmetry imposed | C1 | C1 | C1 |
| Initial particle images (no.) | 2 600 000 | 2 600 000 | 2 600 000 |
| Final particle images (no.) | 18 000 | 104 000 | 16 000 |
| Map resolution (Å) | 3.6 | 3.1 | 3.5 |
| FSC threshold |  |  |  |
| Map resolution range (Å) | 3.3-15 | 2.8-5 | 3.3-9 |
| **Refinement** |  |  |  |
| Initial model used | 8g8g, 6t90 | 8g8g, 6t90 | 8g8g, 6t90 |
| Model resolution (Å) | 3.4 | 3.0 | 3.5 |
| FSC threshold |  |  |  |
| Model resolution range (Å) | 124-3.4 | 124-3.0 | 3.5-124 |
| Model composition |  |  |  |
| Nonhydrogen atoms | 14548 | 12973 | 13981 |
| Protein residues | 977 | 908 | 1035 |
| Nucleotide | 328 | 279 | 279 |
| *B* factors (Å<sup>2</sup>) |  |  |  |
| Protein | 90.31 | 93.38 | 103.14 |
| Nucleotide |  | 97.20 | 132.95 |
| R.m.s. deviations | 213.58 |  |  |
| Bond lengths (Å) |  | 0.004 | 0.005 |
| Bond angles (°) |  | 0.693 | 0.757 |
| Validation |  |  |  |
| MolProbity score | 1.76 | 1.86 | 2.16 |
| Clashscore | 11.31 | 7.49 | 12.56 |
| Poor rotamers (%) | 0.12 | 3.03 | 2.66 |
| Ramachandran plot |  |  |  |
| Favored (%) | 96.85 | 97.63 | 96.54 |
| Allowed (%) | 3.15 | 2.37 | 3.36 |

**Supplementary Table 2.** Cross-correlation between indicated maps and models with DNA at 4 different positions. The highest value is indicated in bold.

| register | 0 | 1 | 2 |
| --- | --- | --- | --- |
| Sox2A | 0.68 | **0.74** | 0.67 |
| Sox2B | 0.59 | **0.68** | 0.61 |
| Sox2C | 0.68 | **0.71** | 0.66 |
| Oct4_Sox2E | 0.72 | **0.75** | 0.72 |
| 2Oct4_Sox2F | 0.68 | **0.75** | 0.68 |

**Supplementary Table 3.**
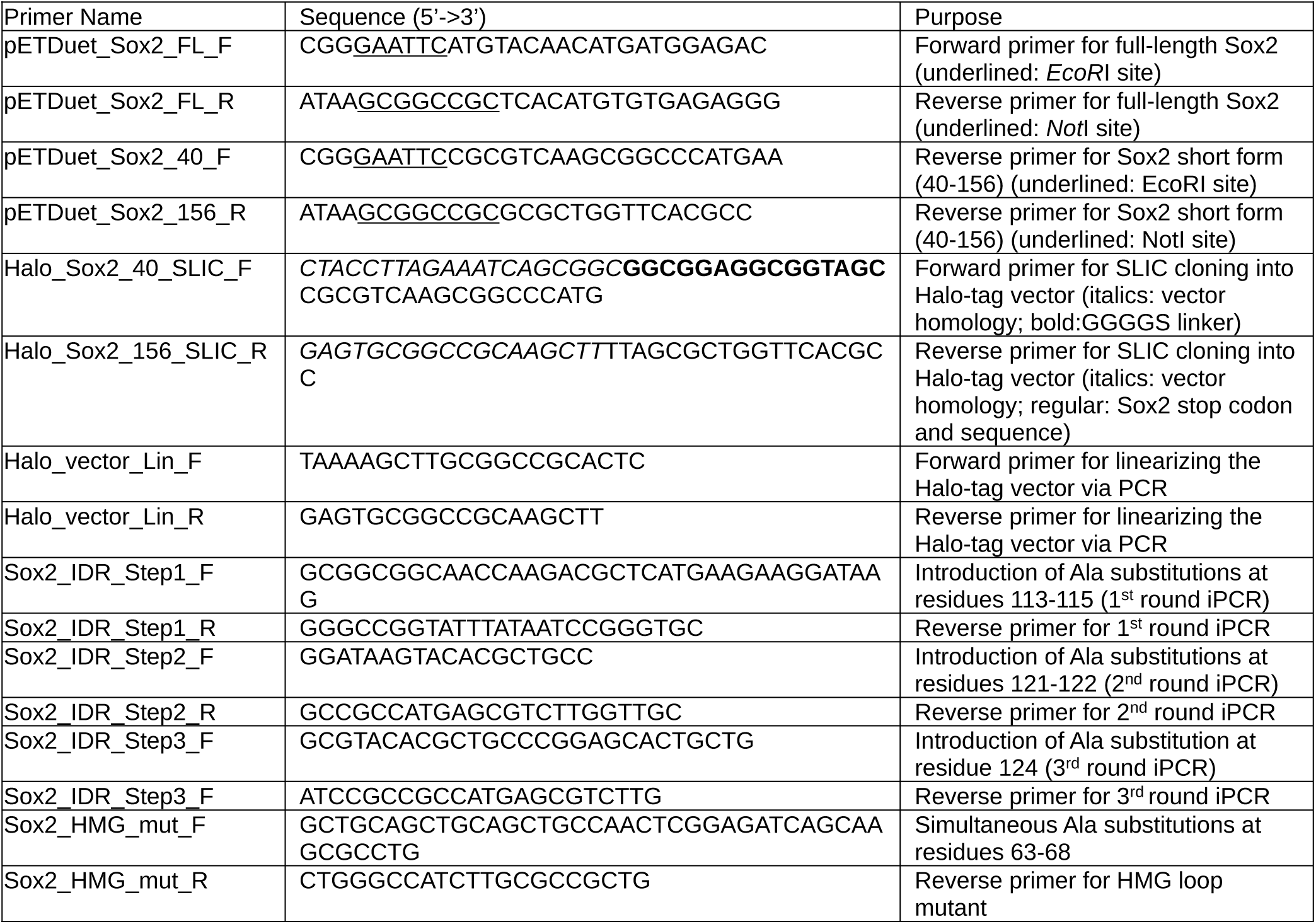
Oligonucleotides used in this study.

**Supplementary Table 4.** Plasmids used in this study.

| Plasmid Name | Backbone | Insert | Tag | Reference |
| --- | --- | --- | --- | --- |
| pETDuet_Sox2_FL | pETDuet-1 | human Sox2 full-length | 6xHis | This study |
| pETDuet_Sox2_S_wt | pETDuet-1 | human Sox2 (residues 40-156) | 6xHis | This study |
| pETDuet_Sox2_S_IDR_6A | pETDuet-1 | human Sox2 (40-156) with Ala substitutions at residues 113-115, 121-122, and 124 | 6xHis | This study |
| pET28_Halo_Sox2_S_wt | pET28a | human Sox2 (residues 40-156) | 6xHis_3xHA_Halo | This study (vector from Addgene #188943) |
| pET28_Halo_Sox2_S_HMG_6A | pET28a | human Sox2 (40-156) with Ala substitutions at residues 63-68 | 6xHis_3xHA_Halo | This study |

